# Open source tools and standardized data in cancer systems biology

**DOI:** 10.1101/244319

**Authors:** Paul Macklin, Samuel H. Friedman, MultiCellDS Project

## Abstract

To reach the full potential of multicellular systems biology, mathematical and computational modelers must pool their efforts to share and curate biophysical measurements, create and combine mathematical models, analyze and visualize model predictions, and validate and refine against shared data. An ecosystem of open source software that reads standardized data is essential. We review the state-of-the-art in open source software and data standards in multicellular systems biology, and point out areas of needed growth to move beyond isolated models to community-driven frameworks that shed light on complex problems in multicellular systems biology.

## I. Introduction

Cancer is a complex multicellular systems problem involving many dynamically coupled processes (e.g., mutations, molecular signaling, substrate transport, selection, tumor-immune interactions) at many scales (molecular, cellular, multicellular, tissue, whole-organ, whole-patient). Thus, numerous modeling approaches are needed to simulate cancer, including molecular-scale models (e.g., systems of ordinary differential equations (ODEs)), cell- and multi-cellular-scale models (usually discrete or agent-based models (ABMs)), continuum-scale biotransport and tissue mechanics (systems of partial differential equations (PDEs)), and multi-systems models (e.g., coarse-grained systems of ODEs). Moreover, these modeling components must be adapted to particular cancer and host physiological processes, and dynamically linked to model the *systems* aspects of cancer [1,2].

To develop these multi-systems cancer models, simulation packages must be able to dynamically share and modify data. This requires a *data model* with a unified representation of biological and physical parameters coming from a well-defined dictionary (an *ontology*). See Fig. 1 (top). Moreover, to ensure that scientific insights gleaned from such simulation frameworks are repeatable, reproducible, and extensible, the underlying software must not only be available in the long term, but also as *open source*, providing the modeling community with the full source code to examine and modify, with unambiguous reuse rights via open source licenses [2,3]. Historically, the trend in computational oncology has been the opposite: singleuse computational models written with *ad hoc*, non-standardized data, in-house visualization and data analysis scripts, and limited or no availability of source code. This has hampered integration of single-process models into systems frameworks, increasing the amount effort required by individual labs to create, calibrate, and use mathematical models in cancer. Moreover, it increases the “lock-in” effect where single labs tend to use a single computational approach for all problems, due to steep development and training costs. This negatively impacts reproducibility and collaborative science [2,3]. Ideally, mathematical oncology as a field should evolve towards an *ecosystem* of open source software that can readily be combined by communicating through open data models. See Fig. 1.

**Figure 1.**
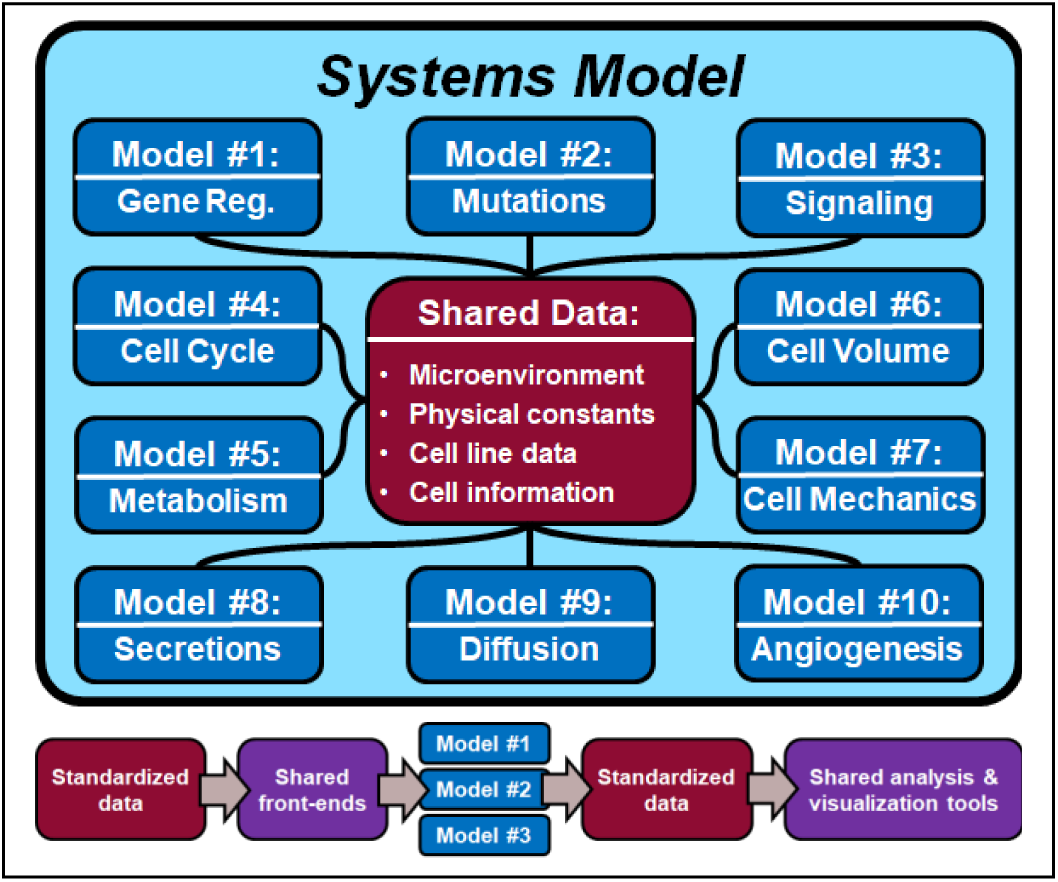
*(top)* Computational models should communicate through a standardized data model, with data elements from well-defined ontologies. *(bottom)* An ecosystem of open source modeling components and tools, connected with open, standardized data, can scale to larger cancer systems problems.

## II. Current Standards and Software

We briefly survey the current state of the field. We cover the most major open source packages used in cancer and related tissue biology, but omit some less-related/used tools.

## A. Standards

*Ontologies:* The greatest progress to date has been in defining ontologies for molecular processes, particularly the Gene Ontology [4], OM for units of measurement [5], and ChEBI chemical entities [6]. More recently, the Cell Behavior Ontology (CBO) [7] emerged to annotate key cellular processes (e.g., division). See [3] for more discussion.

*Data Standards and Markups:* Here there has also been greatest progress at the molecular scale, such as SBML (Systems Biology Markup Language) to describe systems of ODEs in systems biology [8] and PharmML to describe pharmacodynamics data and models [9]. SBML has attempted to extend to spatial and multicellular processes, but progress has been slow due to the complexity of extending the well-established standard. More recently, MultiCellDS (multicellular data standard) emerged to describe multicellular data, centered around *digital cell lines* that annotate microenvironment-dependent phenotype, and *digital snapshots* that record the single-time state of a multicellular system. It is currently working to embed existing standards for molecular-scale processes, with CBO to describe interactions between cells and their phenotypes [3].

## B. Data Repositories

To date, The Cancer Genome Atlas (TCGA) is the greatest repository for data, extending beyond its original design for genomic data to include pathology, radiology, and new molecular data [10]. More recently, the National Cancer Institute has introduced data portals such as the *Genomic Data Commons Data Portal* as integrated frameworks for sharing data and building online software that can operate on it [11]. The Allen Cell Explorer has provided some data and software towards working on cells in general, though not focused on cancer [12]. The Human Cell Atlas aims to integrate multiple scales, but it is not yet publicly available [13].

## C. Open Source Simulation Software

*Molecular Scale:* Open source software (OSS) is in a good state at the molecular level: COPASI can run SBML models as a stand-alone application or as part of a larger simulator [14]. libRoadRunner is fast, optimized package to run SBML models designed to integrate with models [15]. MaBoSS is an open source package for Boolean signaling networks [16].

*Single-Cell and Multicellular Scales:* VirtualCell simulates biotransport and other processes in a single cell with complex geometry [17]. NetLogo [18] and Repast [19] can implement cellular automata models (many cells on a lattice without morphology or mechanics). A healthy variety of (mutually data incompatible) cellular Potts frameworks exist to model cells as pixels on a lattice (including morphology), including CompuCell3D [20], Morpheus (well-integrated with SBML via COPASI) [21], and the Tissue Simulation Toolkit [22]. Several open source frameworks have emerged for off-lattice simulation of 10^5^ or more cells with mechanics, including Chaste [23], PhysiCell [24], and Timothy [25].

*Continuum Scale:* Far fewer open source codes exist for continuum-scale biological modelling. BioFVM [26] was designed to simulate many diffusing chemical substrates for use in systems biology; it can also easily implement common Fisher’s type cancer models. State-of-the-art codes use phase field and mixture models to simulate mixtures of cell types with rigorous mechanics [1], but none are currently open source. While many open source finite element codes are available, few are tailored to biology. FEBio is “open source” (source is available with a non-standard license that is incompatible with Open Source Initiative (OSI) licenses like GPL and BSD), but models written in it can be distributed as open source [27]. SimVascular is an open source package to model flow in static (non-evolving) vascular networks [28].

## D. Open Source Support Tools

Currently no comprehensive, unified framework for visualization and analysis of simulation-generated data exists. Numerous open source frameworks allow users to “roll their own”, including VTK (for 3-D rendering based on geometric primitives) [29], R and Python with their built-in libraries and extensions, and ImageJ for image analysis [30]. However, deeper, standardized data analysis remains a pressing need.

## E. Interfacing Open Models with Open Data

Because SBML has been a well-established standard at the molecular scale, an ecosystem of tools has emerged to interface through its data model (see the discussion above). Many R and Python packages read and write SBML data, and integrated frameworks like EPISIM have emerged to graphically edit multicellular models with SBML components [31]. VirtualCell [17] can directly read microscopy data for its cell geometry. Software such as CellPD has been written to support standardized proliferation and death rate analysis of high-content microscopy data using the MultiCellDS data model [32].

## III. Areas for Future Growth

We need to better connect existing standards to span the molecular, tissue, organism, and population scales. MultiCellDS was designed to embed smaller scale ontologies, but more is needed to connect to organism and population scales.

We need to represent, collect and share not just measurements (data models), but also observations and hypotheses using semantic annotations such as CBO and mathematical model languages such as SBML. In [3], we observed the particular need to record cell phenotypic interaction networks and differentiation networks involving multiple cell types. Here, too, we could link data models like MultiCellDS with mathematical model descriptors and semantic annotations.

PhysiCell recently extended MultiCellDS’ hierarchical phenotype data element to introduce *Cell Definitions*, which bundle phenotypes with user-defined functions to modify phenotype [24]. These functions should be annotated with standardized representations. More work is needed to annotate motility; the Cell Migration Standarisation Organisation (CMSO) and MULTIMOT [33] are actively pursuing this problem

High quality data must be readily available, searchable, and *FAIR* (findable, accessible, interoperable, reusable) [34], thus requiring data aggregation and curation. MultiCellDS has created hand-curated databases of community-contributed data [3], but we need bioinformaticians and library scientists to grow these into user-friendly, community-curated resources.

We need to integrate models at multiple scales to identify the scientific questions that should drive simulation software development. Otherwise, we risk missing connections and slowed development. To run simulations replicably between users, we need shared tools to configure the software, models, and the data analysis pipelines, to track the provenance of each step, and to assemble the results under FAIR principles.

We to map molecular and phenotype data onto mathematical models, and store the results. We propose a library of Cell Definitions, which would extend digital cell lines to include molecular data and lists of observed cell behaviors, and network the cell definitions to represent cell-cell interactions. Once annotated, we need standardized ways to map these onto common paradigms. Lastly, the community must form standards for data quality and validation to drive community curation and quality control.

